# Screening under infection-relevant conditions reveals chemical sensitivity in multidrug resistant invasive non-typhoidal *Salmonella* (iNTS)

**DOI:** 10.1101/2022.09.16.508293

**Authors:** Caressa N. Tsai, Marie-Ange Massicotte, Craig R. MacNair, Jordyn N. Perry, Eric D. Brown, Brian K Coombes

**Affiliations:** Department of Biochemistry and Biomedical Sciences, McMaster University, Hamilton, ON, L8S 4L8, Canada; Michael G. DeGroote Institute for Infectious Disease Research, McMaster University, Hamilton, ON, L8S 4L8, Canada

## Abstract

Bloodstream infections caused by invasive, non-typhoidal salmonellae (iNTS) are a major global health concern. These infections are especially problematic in sub-Saharan Africa, where the sequence type (ST) 313 of invasive non-typhoidal *Salmonella* Typhimurium (iNTS) is dominant. Unlike *S*. Typhimurium strains that cause mild gastroenteritis, iNTS strains are resistant to multiple first-line antibiotics and have higher extraintestinal invasiveness, limiting current treatment options. Here, we performed multiple small molecule screens under infection-relevant conditions to reveal chemical sensitivities in ST313 as entry points to drug discovery to combat the clinical burden of iNTS. By screening the invasive ST313 sequence type under host-mimicking conditions, we identified the antimicrobial activity of the nucleoside analog 3’-azido-3’-deoxythymidine, which required bacterial thymidine kinase activity for its antimicrobial activity. In a parallel macrophage-based screening platform, we also identified three host-directed compounds (amodiaquine, berbamine, and indatraline) that significantly restricted intracellular replication of ST313 in macrophages without directly impacting bacterial viability. This work provides evidence that despite elevated invasiveness and multidrug resistance, iNTS *S*. Typhimurium remains susceptible to unconventional drug discovery approaches.

## INTRODUCTION

*Salmonella enterica* is an important global pathogen that causes disease in a wide range of animal hosts. Pathogenic lifestyles within this species exist along a spectrum, from non-typhoidal serovars that occupy a broad host range and generally cause uncomplicated gastroenteritis, to typhoidal serovars that are host-restricted and linked to bloodstream infection (1). In humans, *Salmonella* Typhimurium commonly causes self-limiting gastroenteritis. Many cases are caused by the ST19 sequence type of non-typhoidal *Salmonella enterica* serovar Typhimurium, which induces inflammation in the gut and proliferates within *Salmonella*-containing vacuoles (SCVs) in epithelial cells and macrophages. In otherwise healthy individuals, ST19 infections are typically confined to the intestinal tract and generally resolve without antibiotic treatment.

In recent years, new clades of non-typhoidal, highly invasive *Salmonella* strains have emerged with infection profiles that more closely resemble typhoidal serovars during human infection (2). These invasive variants can cause lethal septicaemia, clinically termed invasive non-typhoidal *Salmonella* (iNTS) disease, mostly in HIV-positive, malaria-infected, or malnourished individuals (3). iNTS has been most studied in sub-Saharan Africa where a single sequence type, ST313, remains dominant (4). Most ST313 isolates are multidrug resistant to chloramphenicol, ampicillin, kanamycin, streptomycin, sulfonamides, and trimethoprim, rendering most first-line treatment options ineffective and resulting in high case fatality rates (5). Some strains of iNTS ST313 are also resistant to third-generation cephalosporins (6) and azithromycin (7).

The dissemination of antibiotic resistance within iNTS subclades is adding urgency to an already serious problem, as any further restriction in treatment options for iNTS disease is likely to increase global morbidity and mortality. Given pre-existing resistance, addressing this unmet need will require a thorough exploration of alternative therapeutics and screening platforms to identify compounds with activity against ST313 *Salmonella*. Little is currently known about ST313 chemical sensitivity to novel anti-infective agents, as most iNTS studies have been directed to comparisons of ST313 and ST19 infection biology. Indeed, several genomic, phylogenetic, and transcriptomic studies have revealed considerable genomic synteny and collinearity between ST313 and ST19 isolates (8). Comparisons between the two sequence types has identified greater pseudogenization in ST313, particularly in metabolic pathways required for growth in the inflamed gut, consistent with adaptation to extraintestinal pathogenic lifestyles (8) (9, 10). These studies have also revealed increased serum resistance (11), acid tolerance (12), and extraintestinal colonization (13) among ST313 isolates, and decreased swimming motility (14), stationary-phase catalase production (15), biofilm formation (16), epithelial cell invasiveness (17), and induction of macrophage cytotoxicity (15). The impact of these fitness differences on anti-infective activity against ST313 remains unknown, and a systematic study investigating the chemical sensitivity of ST313 in unconventional, infection-relevant growth conditions has not been reported.

In this work, we profiled the chemical sensitivity of ST313 to a broad range of bioactive molecules under infection-relevant conditions. We reasoned that screening in host-mimicking environments may reveal unique vulnerabilities in ST313 that could sidestep existing multidrug resistance, serving as an entry point to guide future anti-iNTS therapeutic development. To this end, we performed two small molecule screens against iNTS ST313 *Salmonella* grown in host-mimicking media and during intracellular infection of macrophages. We first screened a library of 3,840 chemical compounds against ST313 *Salmonella* grown in media that resembles the intracellular SCV, looking to identify compounds with antimicrobial activity in this defined chemical niche. This led to the identification of the nucleoside analog 3’-azido-3’-deoxythymidine (AZT) as an inhibitor of ST313 that was dependent on bacterial thymidine kinase activity. We also screened the same chemical library against ST313 grown in cultured macrophages using a screening design aimed to enrich for host-targeted compounds that enhance host-dependent bacterial killing. This screen identified three compounds (amodiaquine, berbamine, and indatraline) that reduced replication of ST313 *Salmonella* in the intracellular niche.

## RESULTS

### Chemical sensitivity of iNTS ST313 under infection-relevant conditions

Recent studies have illustrated the value of antimicrobial susceptibility testing under infection-relevant growth conditions (18-22), where ionic and nutrient conditions differ from standard bacteriologic media. Considering this, we examined the chemical sensitivity of ST313 *Salmonella* grown in acidic, low phosphate, low magnesium media (LPM), which is known to mimic the chemical and nutrient composition of the intracellular replication vacuole occupied by *Salmonella* in host cells (23). Growth in this host-mimicking media also potentiates antibiotics with normally poor activity against Gram-negative bacteria (21, 24), indicating that host-mimicking conditions can reveal otherwise cryptic antimicrobial activities. Therefore, we screened a chemical library of 3,840 previously approved drugs, natural products, and other biological actives against a representative strain of ST313 iNTS, looking to identify compounds with antimicrobial activity as determined by growth after 20 h of incubation (**Figure 1a**). We also reasoned that systemic, disseminated iNTS disease might result from unique interactions with host immune processes, and by extension, might be susceptible to small molecule manipulation of macrophage defenses. To investigate this possibility, we also screened the same chemical library of 3,840 compounds against ST313 *Salmonella* in a cell-based assay using infected RAW264.7 macrophages. To do this, we designed a screening approach to enrich for compounds with host-targeting activity (**Figure 1b**). Briefly, each compound in the library was added to a well of uninfected RAW264.7 macrophages for 4 h and then washed out prior to macrophage infection with opsonized ST313 *Salmonella* engineered for luciferase expression (25). With slight modifications to our previously reported high-throughput infection method (21), macrophages were treated with fosfomycin following infection to prevent extracellular bacterial replication, and intramacrophage bacterial viability was monitored by reading luminescence immediately after fosfomycin treatment and 6 h later. This screen design limited direct bacterial exposure to library compounds, thereby enriching for host-directed actives.

**Figure 1.**
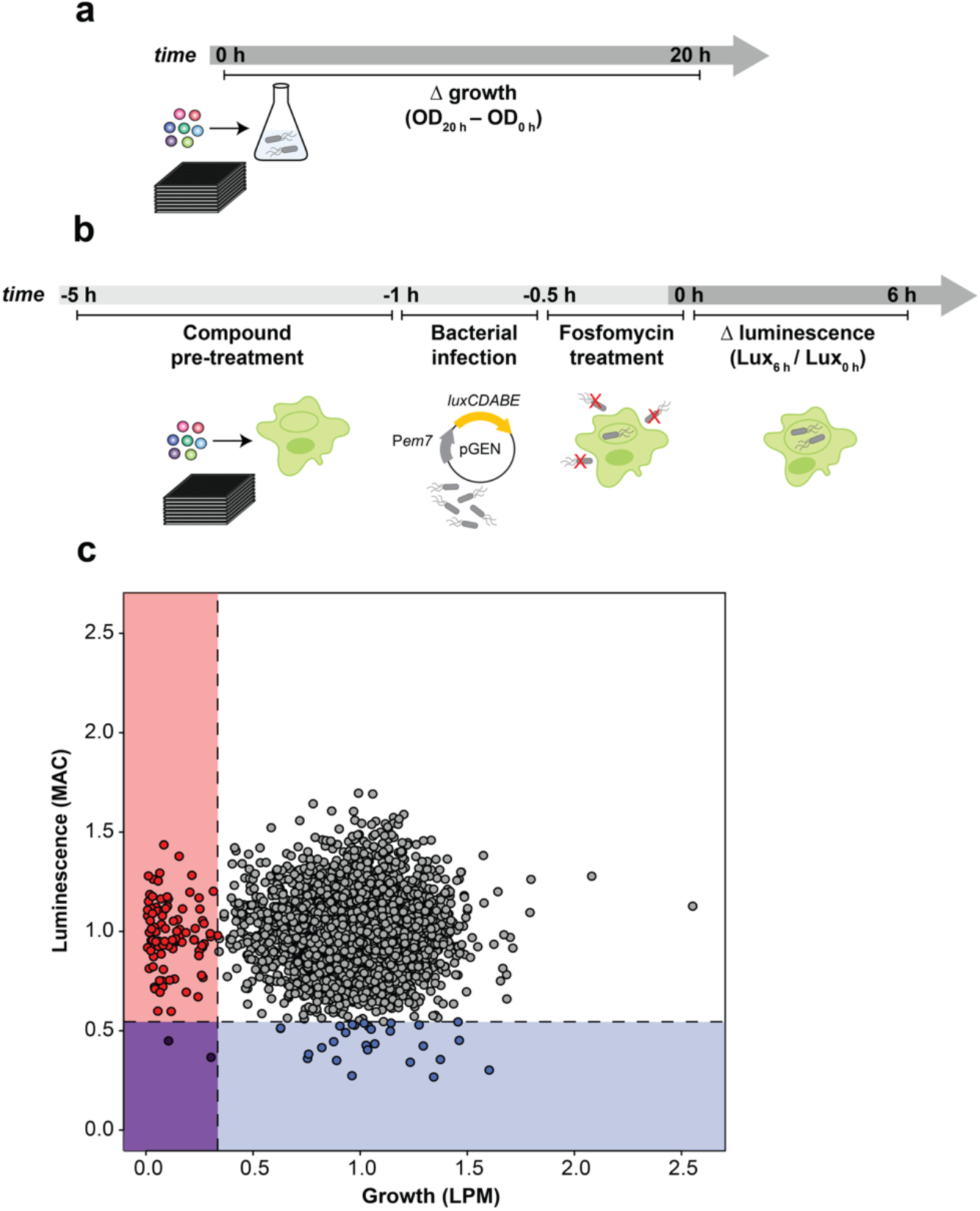
Chemical sensitivity of ST313 grown in infection-relevant conditions. **(a)** Workflow of screening methods for small molecule screen in LPM. OD_600_ was read at 0 h and 20 h after compound addition. **(b)** Workflow of screening methods for small molecule screen in macrophages. RAW264.7 macrophages in 384-well plates were pre-treated with compounds for 4 h, infected with ST313 expressing the luminescent pGEN-*luxCDABE* plasmid for 30 min, then treated with fosfomycin to kill extracellular bacteria for 30 min. Luminescence was read at 0 and 6 h to approximate bacterial viability. **(c)** Plot of screening data from macrophage and LPM chemical screens. 3,840 chemical compounds were tested against ST313 *Salmonella* grown in both conditions, each in technical duplicate. Along x axis, bacterial growth was monitored, values on graph represent interquartile mean-normalized, background subtracted OD_600_ over 20 h of incubation. Along y axis, luminescence production from pGEN-*luxCDABE* was monitored over 6 h, values on graph represent interquartile mean-normalized RLU per well, at 6 h divided by 0 h. Boxes and points indicate compounds that reduced bacterial growth and luminescence to 2.65 s.d. (dotted lines) below the mean of the dataset (blue, luminescence only; red, growth only; purple, both).

After normalizing each screening dataset independently and correcting for plate and well effects by interquartile-mean based methods (26), we directly compared the growth of ST313 over 20 h in LPM (OD_600_) to the replication (luminescence) of ST313 over 6 h in RAW264.7 macrophages after exposure to each chemical compound (**Figure 1c, Dataset S1**). Using a standard deviation cut-off to identify hit significance, we found 92 compounds that restricted ST313 growth in LPM and 30 compounds that restricted ST313 luminescence in macrophages. Only two compounds, cetylbyridinium chloride (and antiseptic quaternary ammonium compound) and purpurogallin carboxylic acid (an oxidation product of gallic acid with anticancer activity) were active in both screens, demonstrating the selectivity of our revised macrophage screening platform to identify host-targeted molecules that elicit antimicrobial activity (21).

### Potency analysis of AZT against ST313

The 92 compounds with antimicrobial activity against ST313 *Salmonella* under host-mimicking conditions included several quinolone antibiotics, antifungals, and other known antimicrobials. Based on potency, chemical diversity, and selectivity, we selected 3’azido-3’deoxythymidine (AZT) for follow-up experiments. AZT is an azido-substituted thymidine analog and antiretroviral agent used in the treatment of HIV. After re-ordering, we determined that AZT inhibited ST313 growth at ∼0.5 μg/mL in LPM media and rich media (**Figure 2a**). For comparison to other pyrimidine analogs, we also tested the fluorinated pyrimidines 5-fluorocytosine (5-FC, **Figure 2b**) and 5-fluorouracil (5-FU, **Figure 2c**) against ST313 in LPM media and rich media.

**Figure 2.**
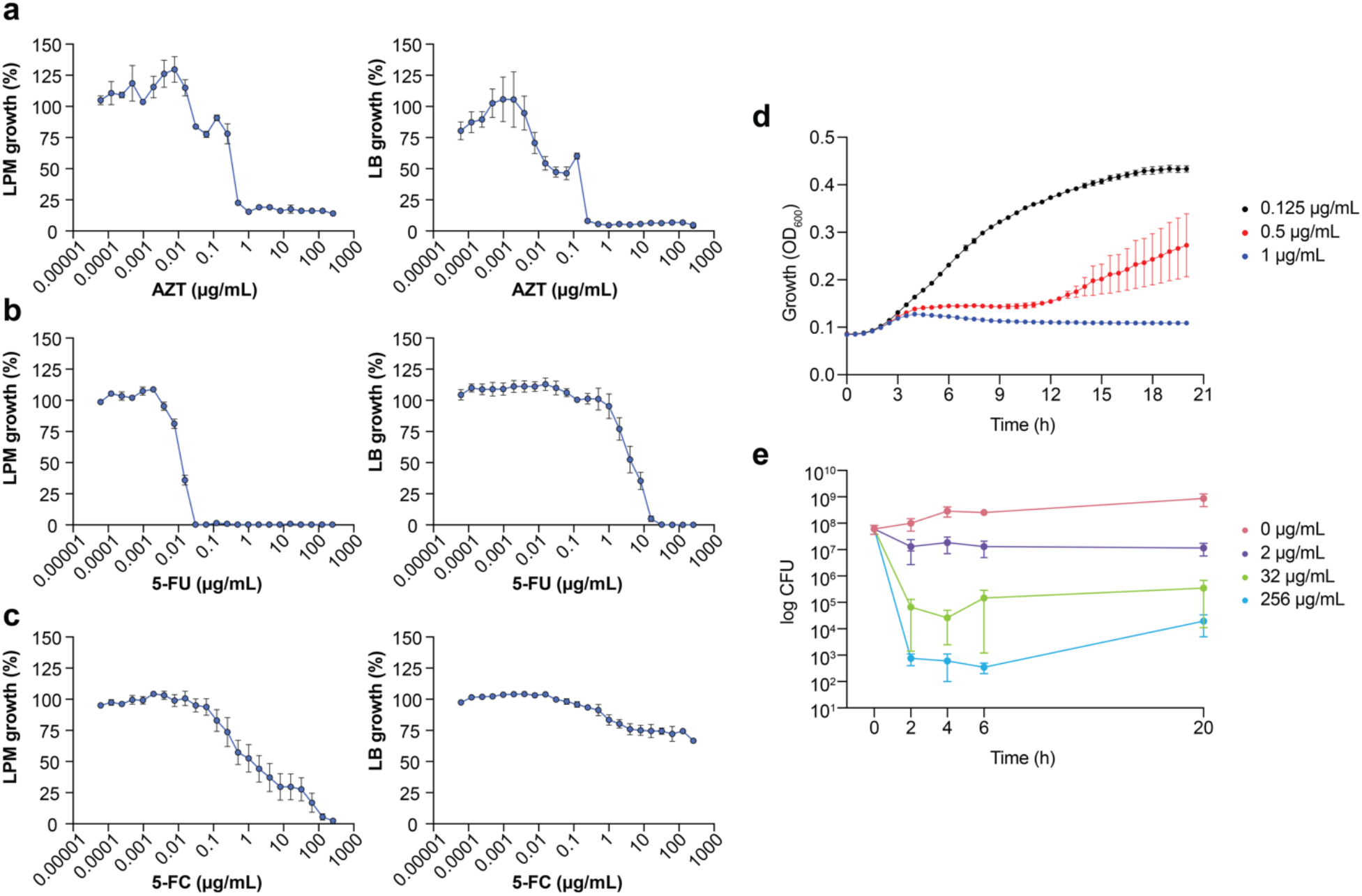
Potency analysis of AZT against ST313. **(a)** Potency of 3’-azido-3’-deoxythymidine (AZT) against ST313 *Salmonella* grown in LPM (left) or LB (right). Growth is normalized to a DMSO control (set to 100%), dots and error indicate mean and s.e.m. for three biological replicates. **(b)** As in panel a, for 5-fluorouracil (5-FU). **(c)** As in panel a, for 5-fluorocytosine (5-FC). **(d)** Growth kinetics of ST313 *Salmonella* in the presence of AZT. OD_600_ was measured at 30 min intervals for 20 h of incubation in LPM, in the presence of AZT at 1 (blue), 0.5 (red), or 0.125 (black) μg/mL. Dots and error indicate mean and s.e.m. for three biological replicates. **(e)** Viable bacterial counts (log CFU) of ST313 *Salmonella* measured after 0, 2, 4, 6, and 20 h of incubation in LPM in the presence of AZT at 256 (blue), 32 (green), 2 (purple) or 0 (pink) μg/mL. Dots and error indicate mean and s.e.m. for three biological replicates.

Interestingly, both 5-FC and 5-FU were active against ST313 in LPM but with different MICs (MIC_5-FC_ ∼ 0.03 μg/mL, MIC_5-FU_ ∼ 100 μg/mL). However, the antimicrobial activity of both analogs was dramatically decreased in rich media, with 5-FC active at only 10 μg/mL in rich media, and 5-FU losing activity entirely in rich media. These data suggested that the antimicrobial activity of AZT towards ST313 is not linked to a conserved aspect of pyrimidine biosynthesis or metabolism.

To gain further insight into the mechanism of AZT activity, we monitored the growth of ST313 in the presence of AZT at sub-inhibitory concentrations during overnight incubation. This approach has been shown to generate unique dose response profiles for different antimicrobial agents, providing insight into mechanisms of bactericidal or bacteriostatic activity (27). In the presence of AZT at near-MIC concentrations (1 μg/mL and below), ST313 iNTS had an increased lag time, reduced replication, and decreased maximum optical density (**Figure 2d**). Similar results were observed by assessing viable bacterial counts of ST313 on solid media after treatment with AZT at 0, 2, 32, and 256 μg/mL. Concentrations at or above 32 μg/mL reduced bacterial viability of ST313 after 6 h of incubation, and after 22 h of incubation there was a ∼4.5-log decrease in ST313 cell viability relative to untreated bacterial cells (**Figure 2e**). Together, these findings are consistent with dose-dependent bactericidal activity of AZT against ST313.

### AZT activity against ST313 iNTS requires bacterial thymidine kinase

To help guide our investigation into the mechanism of AZT-mediated killing, we tested a set of antibiotics covering several major drug classes for changes in MIC when in combination with AZT. Antibiotic partners included DNA-damaging molecules, macrolides, transcription and translation inhibitors, and beta-lactams. These experiments revealed a synergistic interaction between AZT and ciprofloxacin and AZT in combination with colistin against ST313 grown in both LPM (**Figure 3a**) and LB (**Figure 3b**). Ciprofloxacin is a fluoroquinolone antibiotic that inhibits DNA synthesis, suggesting a possible complementary effect with the nucleoside analog activity of AZT. These results are also consistent with a previous report of synergism between AZT and colistin against other Enterobacteriaceae (28).

**Figure 3.**
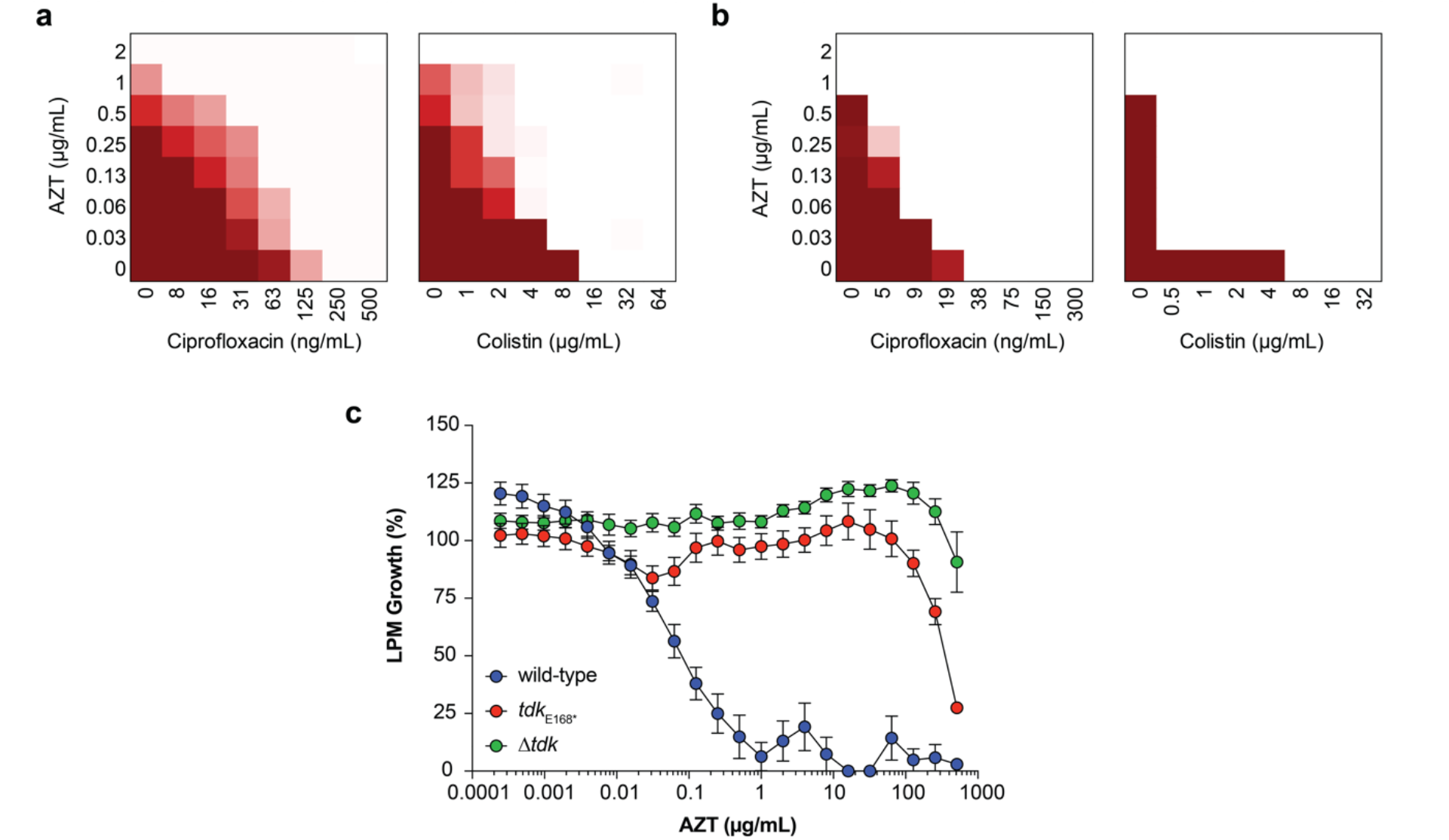
Thymidine kinase-dependent activity of AZT against ST313. **(a)** Checkerboard broth microdilution assays between AZT and ciprofloxacin or colistin, for ST313 grown in LPM. Darker shades of red indicate higher bacterial cell density. **(b)** As in panel a, for bacteria grown in LB. **(c)** Potency of AZT against wild-type, *tdk*_E168*_, and Δ*tdk* ST313 *Salmonella* grown in LPM. Growth is normalized to a DMSO control (set to 100%), dots and error indicate mean and s.e.m. for three biological replicates.

We hypothesized that the antimicrobial activity of AZT was derived from its ability to exploit bacterial metabolism and become incorporated into DNA, and that ST313 susceptibility was attributable to either nucleotide metabolism or processing of AZT itself. To assess this hypothesis, we aimed to isolate mutants that were resistant to AZT. Serial passage of ST313 in the presence of AZT yielded one AZT-resistant strain that exhibited a shift in MIC from 0.5 μg/ml to >512 μg/ml. Whole-genome sequencing of this strain revealed one acquired nonsense mutation (E168*) in a gene encoding for thymidine kinase (*tdk*; STMMW_17451). This finding aligns with previous data indicating AZT and other nucleoside analogs act as prodrugs that require phosphorylation by endogenous kinases. AZT can be phosphorylated by thymidine kinase into its 5’-mono-, -di-, and -triphosphate derivatives (29), and only AZT-triphosphate can be incorporated into growing DNA chains to terminate chain elongation (30). To confirm these results, we transferred the *tdk*_E168*_ nonsense mutation into a wild-type ST313 iNTS background by allelic replacement and also generated a Δ*tdk* mutant and tested the susceptibility of these strains to AZT. Both the *tdk*_E168*_ and Δ*tdk* strains were resistant to AZT activity relative to wild-type ST313, with a >500-fold shift in MIC (**Figure 3c**). These data suggested that AZT activity against ST313 is dependent on bacterial thymidine kinase.

### Potency analysis of macrophage-active compounds

Our host-directed chemical screens identified 28 compounds that significantly restricted intracellular replication of ST313 in cultured macrophages that had been pre-treated with compounds (**Figure 1c**). We presumed that these compounds were host-targeted because they were not hits in the host-mimicking screen performed earlier. To further investigate these putative actives, we re-ordered and re-screened all 28 compounds for potency and found that 8 compounds resulted in a dose-dependent reduction in intramacrophage ST313 luminescence over a 6 h period (**Figure 4a**).

**Figure 4.**
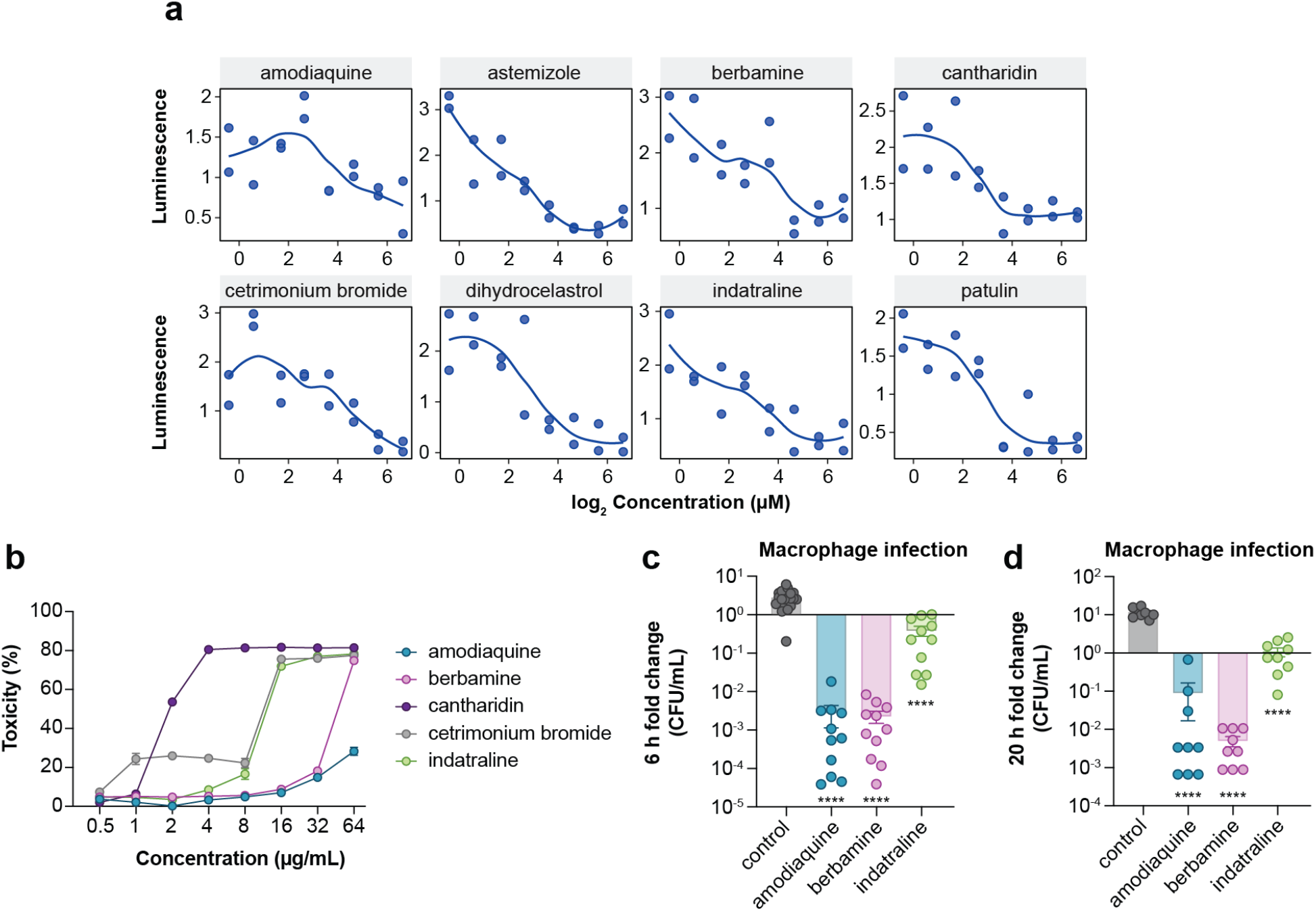
Potency analysis of macrophage-active compounds. **(a)** Re-screening of hit compounds against ST313 *Salmonella* in macrophages. Dots indicate averaged bacterial luminescence from two technical replicates. **(b)** Percentage cytotoxicity in RAW264.7 macrophages after exposure to indicated compounds. Dots indicate mean and s.e.m. from three independent experiments. **(c)** Compound treatment and bacterial infection of RAW264.7 macrophages for 6 h. Bars indicate mean and s.e.m. Groups were compared against control-treated macrophages (equivalent concentration of DMSO) via one-way ANOVA and corrected for multiple comparisons with Sidak’s test. ****p < 0.0001. **(d)** As in panel c, for 20 h of bacterial infection.

Based on known activity, chemical diversity, and commercial availability, five compounds were selected for further characterization (amodiaquine, berbamine, cantharidin, cetrimonium bromide, indatraline). We first determined the cytotoxicity of each compound towards macrophages, reasoning that host cell cytotoxicity could impact bacterial luminescence in a manner unrelated to the host defense systems we were interested in. Thus, we monitored lactate dehydrogenase (LDH) release from uninfected macrophages treated with the five prioritized compounds (**Figure 4b**). From these data, we excluded cantharidin and cetrimonium bromide from further experimentation because both compounds resulted in >20% cytotoxicity at or below 4 μg/mL. We also identified a maximum, non-toxic concentration for the remaining three compounds to be used in secondary assays (32 μg/mL for amodiaquine and berbamine, and 8 μg/mL for indatraline).

We next validated the ability of the remaining three compounds to attenuate ST313 replication by enumerating bacterial CFU rather than monitoring luminescence production. Following a similar infection protocol as was used in our high-throughput macrophage screen, we pre-treated macrophages separately with the three remaining compounds at the maximum concentrations described above, then removed each compound and infected macrophages with iNTS ST313 to enumerate CFU at 0, 6, and 20 h post-infection. These data aligned with our luminescence screening results, as treatment of macrophages with each compound significantly reduced the intracellular replication of ST313 at 6 (**Figure 4c**) and 20 h (**Figure 4d**) post-infection.

### Specificity and immunomodulatory activity of host-directed compounds

Despite our efforts to enrich for host-targeting molecules in our screen, it remained possible that some active compounds may have accumulated within macrophages and directly targeted bacterial viability at later stages of the infection. To examine the specificity of amodiaquine, berbamine, and indatraline against intramacrophage *Salmonella*, we tested these compounds against extracellular iNTS ST313 grown in tissue culture media, reasoning that any compounds that were inactive against *Salmonella* in the same experimental media but without macrophages were likely mediating their effects through macrophage function. At the highest compound concentration tested, only berbamine inhibited bacterial growth to ∼50% relative to untreated bacteria, while amodiaquine and indatraline resulted in ∼0 and 20% growth inhibition, respectively (**Figure 5a**), strongly suggesting that the inhibitory activity of these compounds against intracellular iNTS is directed through macrophage dependent activities.

**Figure 5.**
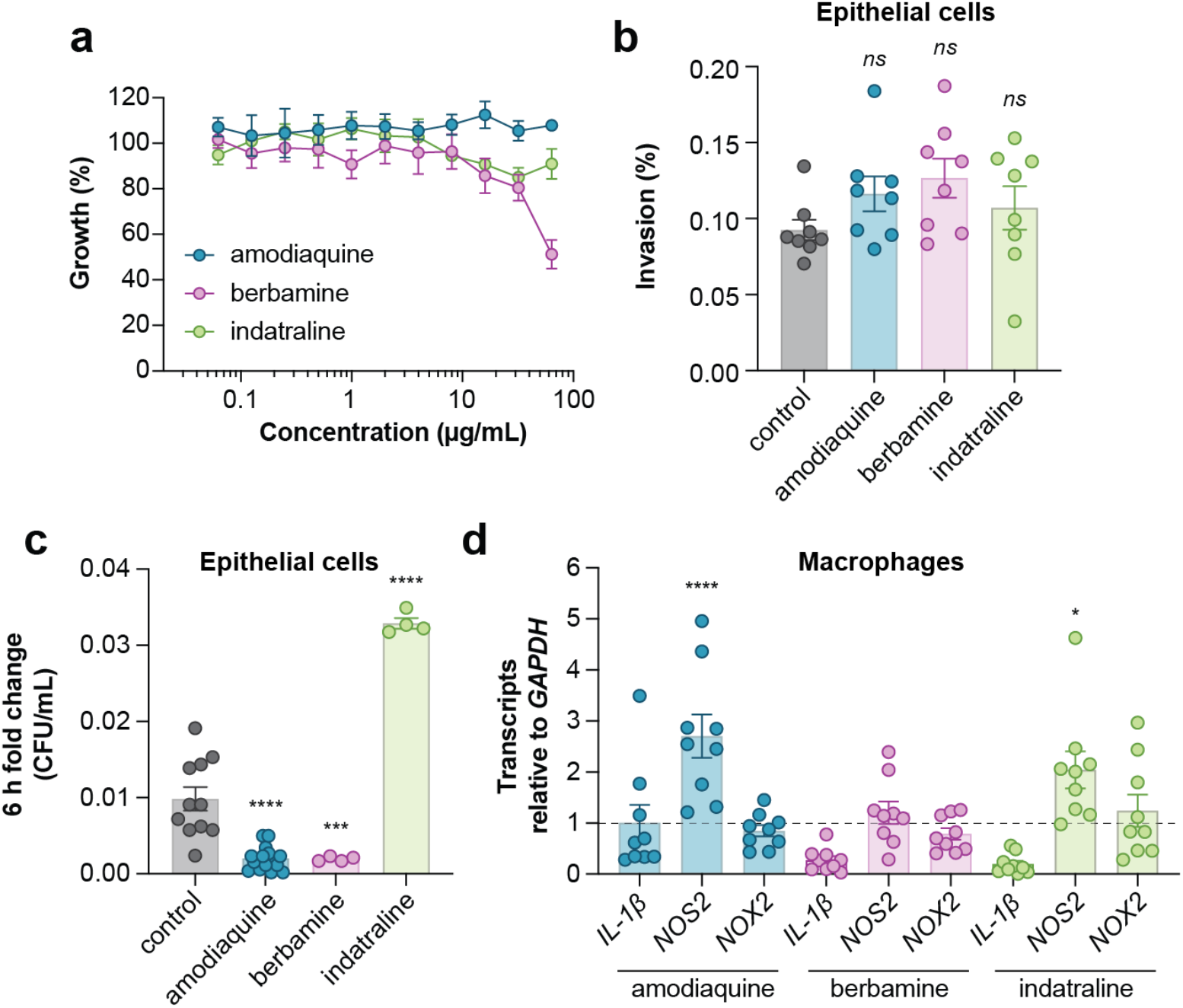
Specificity and immunomodulatory activity of macrophage-active compounds. **(a)** Growth normalized to a DMSO control (set to 100%) of ST313 in the presence of indicated compounds. Data is from four biological replicates, dots and error indicate mean and s.e.m. **(b)** Compound treatment and bacterial infection of HeLa cells. Bars indicate mean and s.e.m. of invasion as a percentage. Groups were compared against control-treated macrophages (equivalent concentration of DMSO) via one-way ANOVA and corrected for multiple comparisons with Dunnett’s test. **(c)** As in panel b, bars indicate mean and s.e.m. of CFU/mL reported as a fold change over 6 h of replication. ***p < 0.001, ****p < 0.0001. **(d)** Gene expression measured by RT-qPCR. Bars indicate mean and s.e.m. Groups were compared against a value of 1 (control-treated macrophages) via one-way ANOVA and corrected for multiple comparisons with Dunnett’s test. *p < 0.05, ****p < 0.0001.

The lack of direct bactericidal activity of amodiaquine, berbamine, and indatraline against iNTS suggested a potential for host-targeting activity that may or may not be specific to macrophages. To examine host cell range, we treated HeLa epithelial cells with these three compounds, infected them with iNTS and monitored bacterial invasion and replication. We speculated that any difference in compound activity between these two cell types might suggest that the compound targeted a macrophage-specific process, such as phagocytosis. In these experiments, none of the compounds inhibited *Salmonella* invasion into epithelial cells (**Figure 5b**), however, like infected macrophages, *Salmonella* replication was significantly inhibited in epithelial cells treated with amodiaquine or berbamine. Unexpected, we found that epithelial cells treated with indatraline supported better *Salmonella* growth as compared to untreated cells (**Figure 5c**). Together, these data suggest that amodiaquine and berbamine likely modulate a host defense pathway that is conserved between epithelial cells and macrophages, while indatraline may impact either distinct pathways in these cell types or the same pathway that functions in different ways to control bacterial growth.

To investigate the possible impact of each compound on macrophage defense functions, we measured the transcription of three key inflammatory-associated genes in RAW 264.7 macrophages including the pro-inflammatory cytokine *IL-1β*, inducible nitric oxide synthase (*NOS2*), and NADPH oxidase (*NOX2*). In these experiments, we observed that both amodiaquine and indatraline significantly increased *NOS2* expression (**Figure 5d**), suggesting that these compounds may promote the production of nitric oxide-dependent host defenses to help clear the intracellular compartment of invasive bacteria.

## DISCUSSION

Bloodstream infections caused by iNTS *Salmonella* are a major health concern. Due to the high prevalence of multidrug resistance in this *Salmonella* pathovar, developing new therapeutics effective against iNTS will require an exploration of unconventional screening approaches and a deeper understanding of chemical sensitivity under infection-relevant growth conditions. In this work, we performed small molecule screens in both nutrient-limited growth media and cultured macrophages to explore the chemical sensitivity of the dominant iNTS serotype, ST313. We identified the antiretroviral nucleoside analog AZT to be active against ST313 in nutrient-limited media in a thymidine kinase-dependent manner. We also discovered three compounds, amodiaquine, berbamine, and indatraline, that significantly restricted the intracellular replication of ST313 in cultured macrophages, likely by augmenting host immune processes and not bacterial viability directly. Preliminary evidence suggests at least two of these compounds may augment nitric oxide-dependent macrophage defense functions to restrict intracellular *Salmonella*, however more work is required to establish a mechanistic understanding of intracellular bacteria growth restriction.

Our first chemical screen performed in host-mimicking media identified AZT as an antibacterial with potent activity against ST313. The antibacterial activity of AZT against other Gram-negative pathogens including *Escherichia, Klebsiella, Shigella*, and *Enterobacter* has been explored. In general, differences in bacterial sensitivity to nucleoside analogs have been linked to variation in the number and substrate specificities of endogenous nucleoside kinases (29)(31). Indeed, our experiments that yielded an AZT-resistant suppressor mutation in *tdk* indicates that bacterial thymidine kinase activity is required for AZT-dependent inhibition in iNTS.

Nucleotide metabolism represents a possible target that should be further explored to identify additional compounds with antimicrobial activity against ST313. A previous transcriptomics study identified expression differences in metabolic genes between ST313 and ST19, including the downregulation of genes involved in uracil and cytosine uptake (*uraB, codB*), melibiose utilization (*melAB*), carbamoyl-phosphate metabolism and pyrimidine biosynthesis (*carAB, pyrEIB*), nitrate reductase (*napDF*), and sulfate metabolism (*cysPU* and *sbp*) (10). ST313 was also previously shown to grow less efficiently than ST19 using purine and pyrimidine as phosphorus sources (32). These studies, in conjunction with our results, indicate that unique transcriptional profiles of ST313 may produce phenotypes susceptible to chemical inhibition. Future experiments directed at understanding the regulation and activity of metabolic and other pathways in iNTS could reveal new entry points into the discovery of additional anti-ST313 compounds.

Previous studies have examined AZT activity in macrophage and murine infection models with *Salmonella* and other Gram-negative bacteria. AZT was shown to reduce intracellular replication of *Salmonella* (of an unknown sequence type) in cultured macrophages after 24 h of incubation (33), and subcutaneous administration of AZT was shown to reduce *Salmonella* burden in infected calves (34). Validating the activity of AZT against ST313 *Salmonella* in both cell culture models of infection as well as *in vivo* remain important steps to continue characterizing the potential of nucleotide metabolism as an anti-ST313 target. Given the synergistic activity we revealed between AZT and colistin or ciprofloxacin *in vitro*, a deeper exploration of combination therapy for ST313 is warranted.

Our second chemical screen was performed in cultured macrophages and was designed to enrich for compounds with host-directed activities. These experiments revealed three compounds (amodiaquine, berbamine, and indatraline) that restricted intracellular replication of ST313 and possibly exert immunomodulatory effects on host cells to increase their innate defense functions. Amodiaquine is an aminoquinoline derivative with anti-malarial and anti-inflammatory properties (35), berbamine is an anti-cancer drug with inhibitory activity towards a *bcr/abl* fusion gene, NF-κB, and IL-1 (36-38), and indatraline is a non-selective monoamine transporter that blocks dopamine, norepinephrine, and serotonin reuptake (39) and is also known to induce autophagy (40). Understanding the full extent of the host processes affected by these compounds requires further experimentation, including transcriptional profiling of macrophages and other cell types.

The likelihood that these compounds modulate host and not bacterial functions would categorize them as host-directed therapies, which comprise chemical agents that either enhance or blunt inflammatory processes activated by the immune system (41). Apart from some studies focusing on host-directed therapies as treatments for *Mycobacterium tuberculosis* infection (41), host-directed therapies have been relatively underexplored as adjuncts to anti-infective therapy. The success and promise of host-directed therapies to modulate innate immune functions in other therapeutic areas (42) (43) warrants a critical exploration of their potential against drug-resistant infections such as those caused by iNTS. The work presented here provides a high-throughput platform and screening approach to probe even greater chemical space towards this goal.

Overall, our findings add to a growing body of work characterizing the fitness of ST313 *Salmonella* under host-relevant conditions and revealed novel chemical sensitivity in iNTS when screened under unconventional approaches. Our data show that infection-relevant growth conditions expose new bacterial vulnerabilities that can be exploited by small molecule targeting, whether through direct antimicrobial action or by targeting host pathways. Larger chemical libraries used in conjunction with the screening approaches described will allow for a more comprehensive chemical susceptibility profile of ST313 *Salmonella*. We consider these results to be encouraging for future therapeutic development against ST313 to overcome existing and widespread multidrug resistance in iNTS.

## METHODS

### Bacterial strains and culture conditions

*Salmonella* experiments were performed with strain D23580 (ST313). For compound screening in macrophages and secondary assays, this strain was transformed with pGEN-*lux* conferring gentamicin resistance (25). Routine propagation of bacteria was in LB media (10 g/L NaCl, 10 g/L Tryptone, 5 g/L yeast extract) supplemented with appropriate antibiotics (streptomycin, 100 μg/ml, gentamicin, 15 μg/ml). Where indicated, bacteria were grown in LPM (acidic pH low Mg^2+^ media) (5 mM KCl, 7.5 mM (NH_4_)_2_SO_4_, 0.5 mM K_2_SO_4_, 80 mM MES pH 5.8, 0.1% casamino acids, 0.3% (v/v) glycerol, 24 μM MgCl_2_, 337 μM PO_43-_). Bacteria were grown at 37°C.

### Cell culture maintenance

Cells were maintained in a humidified incubator at 37°C with 5% CO_2_. HeLa epithelial cells and RAW264.7 macrophages were grown in DMEM containing 10% FBS (Gibco) and seeded in tissue culture-treated 96-well (100 μL/well, 10^5^ cells/well) or 384-well (50 μL/well, 5 × 10^4^ cells/well) plates (Corning) ∼20-24 h prior to use. In experiments with RAW264.7 macrophages, cells were pretreated with 100 ng/mL LPS from *Salmonella enterica* serovar Minnesota R595 (Millipore) for ∼20-24 h prior to infection.

### Reagents

All high-throughput compound screening was performed at the Centre for Microbial Chemical Biology (McMaster University). The chemical library we screened contained 3,840 diverse small molecules assorted from Sigma-Aldrich and MicroSource; screening stocks (5 mM) were stored at −20°C in DMSO. The following compounds were ordered from Sigma-Aldrich: 3’-azido-3’-deoxythymidine (AZT), 5-fluorouracil (5-FU), 5-flurocytosine (5-FC), ciprofloxacin, colistin, amodiaquine, cantharidin, indatraline. Other compounds were sourced from Cedarlane: berbamine, and cetrimonium bromide. Compounds were routinely dissolved in DMSO at a concentration of 10 mg/mL and stored at −20°C.

### LPM chemical screening

Overnight cultures of ST313 *Salmonella* were sub-cultured ∼1:50 in LB and grown for 2-2.5 h. The cultures were diluted ∼1:350 into LPM and dispensed into 384-well black, clear flat bottom (Corning) plates to a final volume of 30 μL per well. 60 nL of each compound (5 mM stocks) was added using an Echo 550 Liquid Handler directly into wells for a final concentration of 10 μM compound per well. OD_600_ was read immediately after compound addition (OD_0 h_) and again after ∼20 h of incubation at 37°C (OD_20 h_). Normalized growth was calculated by subtracting OD_0 h_ from OD_20 h_, then correcting for plate and well effects by interquartile-mean based methods (26). Compounds that reduced growth more than 2.65 s.d. below the mean of the dataset were considered actives. Screening was performed in duplicate.

### MIC determination for nucleoside analogs

ST313 cultures were grown overnight in LB, then diluted ∼1:10,000 into LPM or LB. AZT, 5-FU, and 5-FC were serially diluted two-fold starting at 128 μg/mL to a final concentration of < 0.0001 μg/mL, then added to bacteria-containing media. OD_600_ was read immediately after compound addition (OD_0 h_) then again after ∼20 h of incubation at 37°C (OD_20 h_). Percentage growth was calculated by subtracting OD_0 h_ from OD_20 h_, then normalizing values to a DMSO control (set to 100%).

### Monitoring kinetics of AZT-induced bacterial death

ST313 cultures were grown overnight in LB, then diluted ∼1:500 into LPM. AZT was added to bacteria-containing media at 1, 0.5, and 0.125 μg/mL. OD_600_ was read immediately after compound addition, then every 30 min for 20 h of incubation at 37°C while shaking. At 0, 2, 4, 6, and 20 h, cultures were serially diluted and plated on LB agar to enumerate viable bacterial CFU.

### Checkerboard broth microdilution assays

8×8 matrices of compound were created in 96-well plates (Corning) with two-fold serial dilutions of AZT and various partner antibiotics at 8 concentrations. After overnight growth in LB, bacteria were diluted ∼1:5000 into LPM or LB and added to each well of the 8×8 matrix. After addition of bacteria, plates were incubated at 37°C for ∼20-22 h, before and after which OD_600_ was measured. At least two biological replicates were performed for each assay.

### Suppressor isolation and whole-genome sequencing

Spontaneous resistant mutants to AZT were selected for by serial passage in liquid culture. An ST313 culture was grown overnight in LPM at 37°C, then diluted ∼1:2000 into 200 μL LPM per well in 96-well plates (Corning) containing two-fold serial dilutions of AZT beginning at 256 μg/mL, and incubated overnight at 37°C. The susceptibility of this strain to AZT was then repeatedly tested for several days: every other day, the highest concentration well with observable growth was sub-cultured and grown overnight at 37°C in LPM containing the corresponding AZT concentration at which growth was observed. When the strain displayed resistance to > 256 μg/mL AZT, genomic DNA was extracted using the QIAamp DNA mini kit (Qiagen). Samples were sequenced on a MiSeq 2×250 platform with paired-end reads. Raw reads were processed with FastQC and trimmed with Cutadapt (44) to remove Nextera transposase sequences. Sequencing data was aligned against the reference genome for *Salmonella* ST313 (FN424405) and analyzed using breseq (45) in polymorphism mode with default settings.

### Cloning and mutant generation

All DNA manipulation procedures followed standard molecular biology protocols. Primers were synthesized by Sigma-Aldrich. PCRs were performed with Phusion, Phire II, or Taq DNA polymerases (ThermoFisher). All deletions were confirmed by PCR and verified by DNA sequencing performed by Genewiz Incorporated. An unmarked, in-frame gene deletion mutant of the *tdk* thymidine kinase gene (STMMW_17451) was generated via homologous recombination from a suicide plasmid as described previously (46). Briefly, ∼500 bp upstream and downstream of the target gene were PCR-amplified and spliced together by overlap-extension PCR, using the following primers: Δ*tdk* upstream F (5’-GGGGAGCTCAGCTGGGTATTCCTAAGTCTATCC-3’), Δ*tdk* upstream R (5’-TACCTGAGGTAAAGAGCGGCTTAT-3’), Δ*tdk* downstream F (5’-TTGGTCGCAGGACCTCACCTGA-3’), Δ*tdk* downstream R (5’-gggGGTACCGGGGGATACTCACCGTCTGTCGCT-3’). The resulting deletion allele was digested with KpnI/SacI, ligated into the KpnI/SacI digested suicide plasmid pRE112 (Edwards et al., 1998), and recovered in *E. coli* DH5α λ pir. The sequence-verified construct was then transformed into *E. coli* SM10λ *pir* to create a donor strain for conjugation, and then introduced into wild-type ST313 *Salmonella* via conjugal transfer. Merodiploid clones were first selected on streptomycin and gentamicin, followed by selection for mutants using SacB-based counterselection on 10% (w/v) sucrose and growth at 30°C(47). Similar methods were used for generation of the *tdk*_E168*_ mutant, introducing the point mutation by overlap-extension PCR and chromosomal replacement by allelic exchange, using the following primers: *tdk*_E168*_ upstream F (5’-GGGGAGCTCCTGTCTGAAGATGCCTTCGATGAC-3’), *tdk*_E168*_ upstream R (5’-CCTTGATCAGGACGGCAGGCCTTATAACAGAGGCGAACAGGTGGT-3’), *tdk*_E168*_ downstream F (5’-GTTCATTCCCGCCAATAACCACCTGTTCGCCTCTGTTATAAGGC-3’), *tdk*_E168*_ downstream R (5’-GGGGGTACCCAATGAATGCGGGTAAGTCGACTGC-3’).

### Macrophage chemical screening

100 nL of each compound (5 mM stocks) was added (Echo 550 Liquid Handler) directly to 384-well black, clear flat-bottom plates (Corning) containing LPS-pretreated RAW264.7 macrophages in 50 μL DMEM + 10% FBS, for a final concentration of 10 μM compound per well. Macrophages were incubated with compounds for 4 h at 37°C with 5% CO_2_. Cultures of ST313 expressing pGEN-*lux* were grown overnight in LB with 15 μg/mL gentamicin. 30 min prior to infection (∼3.5 h after compound addition), bacteria were opsonized for 30 min in 20% human serum (Innovative Research) in PBS at 37°C. Compound-containing media was then removed from macrophages and replaced with 50 μL/well of opsonized bacteria (diluted in DMEM + 10% FBS to achieve an MOI of 50:1). Plates were centrifuged at 200 x *g* for 3 min, then incubated for 30 min at 37°C with 5% CO_2_. Bacteria-containing media was then removed from macrophages and replaced with 50 μL/well of DMEM + 10% FBS + 100 μg/mL fosfomycin to eliminate extracellular bacteria. Plates were incubated again for 30 min at 37°C with 5% CO_2_.

Fosfomycin-containing media was then removed from macrophages and replaced with 50 μL/well of DMEM + 10% FBS + 10 μg/mL fosfomycin. Luminescence was read immediately after this media replacement step (Lux_0 h_), plates were incubated for 6 h at 37°C with 5% CO_2_, then luminescence was measured a second time (Lux_6 h_). Normalized luminescence was calculated by dividing Lux_6 h_ by Lux_0 h_ (to represent fold change replication over the course of the experiment), then correcting for plate and well effects by interquartile-mean based methods (26). Compounds that reduced growth to 50% or less than the mean of the dataset were considered actives. Screening was performed in duplicate. For secondary screening of hit compounds, an identical protocol to the initial macrophage screen was followed, with the exception that compounds were serially diluted two-fold starting at 100 μM to a final concentration of 0.78 μM in DMEM + 10% FBS prior to the 4 h incubation period with macrophages.

### Cytotoxicity assays

LPS-pretreated RAW264.7 macrophages were seeded into 96-well plates in DMEM + 10% FBS as described above. Compounds were serially diluted two-fold starting at 64 μg/mL to a final concentration of 0.5 μg/mL, then added directly to macrophages. After 4 h of incubation with compounds at 37°C with 5% CO_2_, plates were centrifuged at 500 x *g* for 2 min and culture supernatant was collected for quantification of lactate dehydrogenase (LDH) release.

Cytotoxicity was quantified colorimetrically (G-biosciences Cytoscan™-LDH Cytotoxicity Assay) wherein LDH activity is measured by recording A_490_ after 20 min incubation with substrate mix at 37°C. Lysis control wells were treated with 10X lysis buffer for 45 min prior to supernatant collection. Percent cytotoxicity was calculated with the formula:

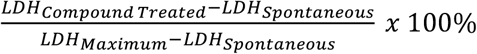 where LDH is the amount of LDH_Spontaneous_ activity in the supernatant of untreated cells and LDH_Maximum_ is the amount of LDH activity in the supernatant of lysis control wells. The LDH activity in cell-free culture medium was subtracted from each value prior to normalization to account for serum.

### Intramacrophage CFU enumeration

LPS-pretreated RAW264.7 macrophages were seeded into 96-well plates in DMEM + 10% FBS as described above. A single concentration of each compound (determined based on cytotoxicity testing) was added directly to wells, with an equivalent volume of DMSO added to control wells. Macrophages were incubated with compounds for 4 h at 37°C with 5% CO_2_. Cultures of ST313 were grown overnight in LB, then 30 min prior to infection (∼3.5 h after compound addition), bacteria were opsonized for 30 min in 20% human serum (Innovative Research) in PBS at 37°C. Compound-containing media was then removed from macrophages and replaced with 100 μL/well of opsonized bacteria (diluted in DMEM + 10% FBS to achieve an MOI of 50:1). Plates were centrifuged at 200 x *g* for 3 min, then incubated for 30 min at 37°C with 5% CO_2_. Bacteria-containing media was then removed from macrophages and replaced with 100 μL/well of DMEM + 10% FBS + 100 μg/mL fosfomycin to eliminate extracellular bacteria. Plates were incubated again for 30 min at 37°C with 5% CO_2_. Fosfomycin-containing media was then removed from macrophages and replaced with 100 μL/well of DMEM + 10% FBS + 10 μg/mL fosfomycin. Immediately after this media replacement step, adhered macrophages from half of the wells were lysed in sterile water. Bacterial CFU from each lysed well were enumerated by serially diluting in PBS and plating on LB agar (CFU at 0 h). After 6 h of incubation at 37°C with 5% CO_2_, adhered macrophages from the other half of the wells were lysed in sterile water for plating and CFU enumeration. Fold change in CFU/mL was calculated (CFU at 6 h divided by at 0 h) to represent replication over the course of the experiment. For experiments with 20 h of incubation, an identical protocol was followed, except for a modified MOI of 20:1 to prevent excessive macrophage lysis.

### MIC determination for amodiaquine, berbamine, indatraline

A culture of ST313 was grown overnight in LB, then diluted ∼1:10000 into DMEM + 10% FBS. Compounds were serially diluted two-fold starting at 64 μg/mL to a final concentration of < 0.1 μg/mL, then added to bacteria-containing media. OD_600_ was read immediately after compound addition (OD_0 h_) then again after ∼20 h of incubation at 37°C (OD_20 h_). Percentage growth was calculated by subtracting OD_0 h_ from OD_20 h_, then normalizing values to a DMSO control (set to 100%).

### HeLa epithelial cell infections and compound treatment

HeLa epithelial cells were seeded into 96-well plates in DMEM + 10% FBS as described above. A single concentration of each compound (determined based on cytotoxicity testing) was added directly to wells, with an equivalent volume of DMSO added to control wells. HeLa cells were incubated with compounds for 4 h at 37°C with 5% CO_2_. A culture of ST313 was grown overnight in LB, then sub-cultured ∼1:50 for ∼2.5 h (beginning ∼1.5 h after compound addition) at 37°C. Compound-containing media was then removed from HeLa cells and replaced with 100 μL/well of bacteria (diluted in DMEM + 10% FBS to achieve an MOI of 100:1). Plates were centrifuged at 500 x *g* for 2 min, then incubated for 10 min at 37°C with 5% CO_2_. Bacteria diluted to the appropriate MOI were serially diluted in PBS and plated on LB agar to enumerate CFU (CFU_input_). Bacteria-containing media was then removed, plates were washed 3 times with PBS, then media was replaced with 100 μL/well of DMEM + 10% FBS and plates were incubated for 20 min at 37°C with 5% CO_2_. Media was then removed and replaced with 100 μL/well of DMEM + 10% FBS + 100 μg/mL fosfomycin to eliminate extracellular bacteria. Plates were incubated again for 30 min at 37°C with 5% CO_2_. Fosfomycin-containing media was then removed, plates were washed once with PBS, then media was replaced with 100 μL/well of DMEM + 10% FBS + 10 μg/mL fosfomycin. Immediately after this media replacement step, adhered HeLa cells from half of the wells were lysed in PBS containing 1% (v/v) Triton-X100 and 0.1% (w/v) SDS. Bacterial CFU from each lysed well were enumerated by serially diluting in PBS and plating on LB agar (CFU_0 h_). After 6 h of incubation at 37°C with 5% CO_2_, adhered HeLa cells from the other half of the wells were lysed in PBS containing 1% (v/v) Triton-X100 and 0.1% (w/v) SDS for plating and CFU enumeration (CFU_6 h_). Percent invasion was quantified by dividing CFU_0 h_ by CFU_input_; fold change in CFU/mL was calculated by dividing CFU_6 h_ by CFU_0 h_.

### RNA isolation and RT-qPCR

LPS-pretreated RAW264.7 macrophages were seeded into 96-well plates in DMEM + 10% FBS as described above. A single concentration of each compound (determined based on cytotoxicity testing) was added directly to wells with 3 technical replicates, with an equivalent volume of DMSO added to control wells. Macrophages were incubated with compounds for 4 h at 37°C with 5% CO_2_. Compound-containing media was removed, then adhered macrophages were scraped and resuspended in 100 μL Trizol (Invitrogen) for cell lysis. RNA was extracted by chloroform (BioShop) separation following the manufacturer’s protocol, precipitated with 100% isopropanol (BioShop) and washed with 75% ethanol (Sigma) before treatment with Dnase I (Turbo DNA-free kit). DNase I was inactivated with 2.5 mM EDTA and RNA was resuspended in DEPC water. For RT-qPCR experiments, cDNA was synthesized from purified RNA using qScript cDNA Supermix (Quantabio) and diluted 1:10 before use. *GAPDH* was used for normalization, RT-qPCR was performed in a LightCycler 480 (Roche) with PerfeCTa SYBR Green Supermix (Quantabio). For all experiments, normalized ratios (compound/DMSO) were calculated relative to *GAPDH* transcript levels.

### Quantification and statistical analysis

Data were analyzed using RStudio version 1.0.143 with R version 3.2.2, and GraphPad Prism 8.0 software (GraphPad Inc., San Diego, CA). Each figure legend contains information on the type of statistical test used as well as mean and dispersion measures. P values of < 0.05 were considered significant.

## Acknowledgements

We thank the McMaster Centre for Microbial Chemical Biology for assistance with high-throughput screening and the Farncombe Metagenomic Facility for technical support with genome sequencing. This work was supported by a research grant to B.K.C. from the Canadian Institutes of Health Research (PJT-156361). C.N.T., M-A.M., C.R.M., and J.N.P. were supported by Canada Graduate Scholarship from the Canadian Institutes of Health Research.

